# Invasive *Argemone mexicana*’s suppressive effects on *Phaseolus vulgaris* and *Zea mays* germination and growth

**DOI:** 10.1101/2023.07.21.550054

**Authors:** Fredrick Ojija

**Affiliations:** Department of Earth Sciences, Mbeya University of Science and Technology, P.O. Box 131, Mbeya, Tanzania

**Keywords:** Allelochemicals, Bioactivity, Allelopathy, Biodiversity, Community ecology, Invasive, Weeds

## Abstract

Invasive *Argemone mexicana* plant species is invading many ecosystems in East Africa. However, there have not been many studies to assess how it affects plants. In petri dishes and pot experiments, we investigated the suppressive effects of *A. mexicana* on *Phaseolus vulgaris* and Zea mays germination and seedling growth. To investigate its suppressive effects on the test plant, different concentrations of *A. mexicana* leaf (AmL) crude extract were applied to the seeds and seedlings of *P. vulgaris* and *Z. mays*. At higher concentrations (70% and 100%), the findings showed that AmL crude extract concentrations reduced the germination and growth of *P. vulgaris* and *Z. mays* seeds. Compared to seeds that germinated at lower concentrations and in the control (0%) group, fewer seeds at higher concentrations grew. Accordingly, higher concentrations, relative to lower ones and controls, retarded seed germination. Additionally, the fresh biomass, root lengths, stem diameters, and heights of *P. vulgaris* seedlings were reduced under 75% and 100% AmL concentrations, which had a negative impact on their growth vigor. Although this study shows that *P. vulgaris* and *Z. mays* germination and growth were inhibited by *A. mexicana* crude extract, field research experiments are needed to investigate the suppressive effects of this invasive weed on other plant species. Due to its detrimental impact on plant growth, the study recommends further management of *A. mexicana* to protect biodiversity. It is expected that these results will be helpful in developing policies and programs for managing invasive plants while taking into account the effects on people’s livelihoods.

## Introduction

Invasive plants, defined here as non–indigenous plants that disrupt systems where they exist, are having a deleterious impact on the environment worldwide [1–3]. They interfere and compete for resources such as nutrients, water, light, and space, including pollination services with native plants as well as crops [4–6]. Natural ecosystems are the most vulnerable to biological invasions of alien invasive plant species (AIP), which alter patterns of biodiversity and ecosystem functioning [4, 7, 8]. In their non– native range, AIPs jeopardize ecological integrity [9–11], food production [3, 12], and the provision of ecosystem services essential for human sustainability [13, 14]. Billions of dollars are spent annually on the monitoring and control efforts of AIPs, which also cause higher food prices and lower farm incomes globally [9, 15, 16].

In addition to being able to compete with and suppress native plants and crops, most AIPs also possess anti–herbivorous and anti–microbial traits, which means they have no natural enemies in their new range [17, 18]. Furthermore, they possess allelochemicals that prevent neighbouring plants from germinating, growing, and/or developing [13, 19]. The climate change has also been acknowledged as the primary driver of AIP’s occurrence dynamics [17]. Because of these traits, some AIPs are able to displace native plant species while dominating invading habitats and/or rangelands [17, 19, 20]. In general, the majority of AIPs and their associated problems significantly impact global economic growth [3, 6, 14, 21, 22].

One of the damaging AIPs that endangers biodiversity and food production in sub–Saharan Africa is the Mexican poppy (*Argemone mexicana* L., Papaveraceae) [23, 24]. It is an invasive herb (with prickles) that can reach a height of 1 m [25–28]. Its yellow, scentless flowers have a diameter of about 4 to 5 cm, and its black, spherical seeds are around 5 to 11 cm long and spiny [25]. Fig 1 shows the seedlings, mature or adult plant, flower, and fruit of an invasive *A. mexicana*. Additionally, *A. mexicana* has a 3 cm–long, spiny, obovate capsule [25]. Although *A. mexicana* is indigenous to tropical America (e.g., Mexico, the United States, the Virgin Islands, India, and Nicaragua), it is an invasive species in other countries including Botswana, Tanzania, Zimbabwe, Côte d’Ivoire, Mauritius, Cuba, Syria, and Kansas [23, 29, 30]. Also, *A. mexicana* has been naturalized in some countries, and it is thus considered an agricultural weed [31].

**Fig. 1.**
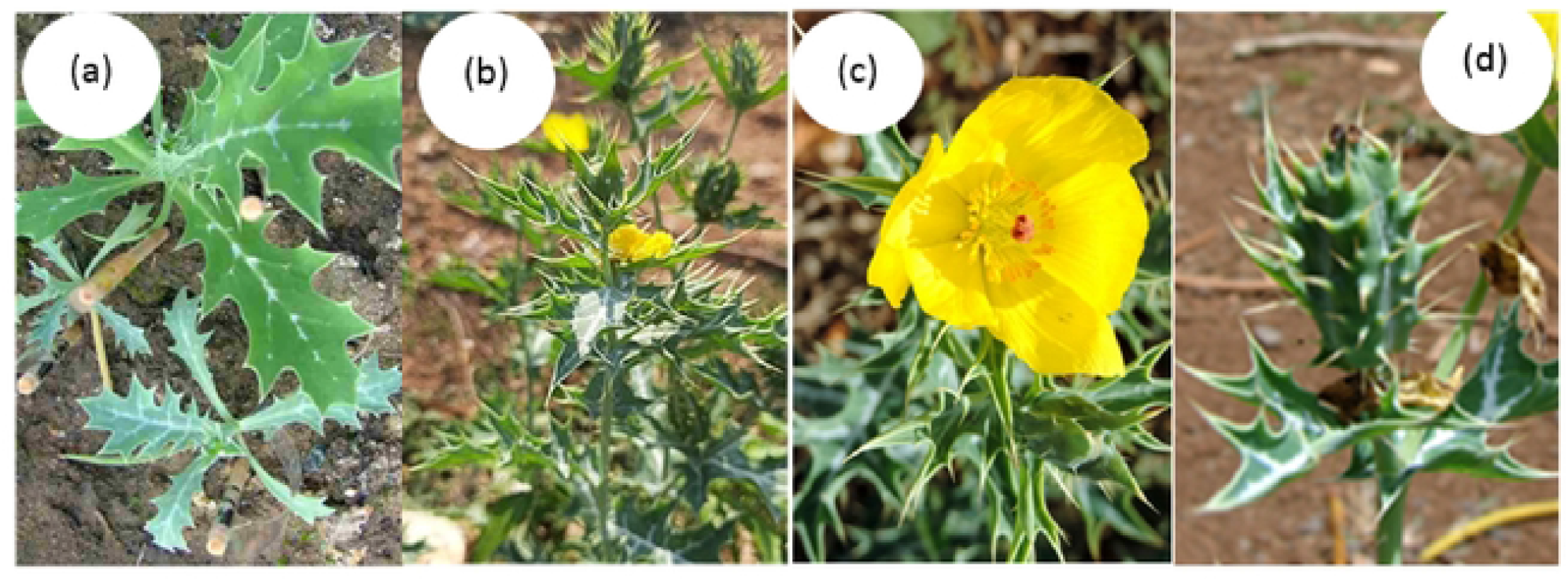
The (a) seedlings, (b) mature or adult plant, (c) flower, and fruit of an invasive *Argemone mexicana*. Photos: F. Ojija, 2022.

Like many other IAPs, *A. mexicana* can colonize a range of habitats, including savannas and grasslands. These include both disturbed and undisturbed environments [23]. Agricultural areas, waster areas, pasturelands, construction sites, floodplains, and along road verges or roadsides are examples of areas where *A. mexicana* frequently infiltrates [25]. Moreover, it has been reported that it can also invade arable land and rangelands [23, 30].

According to Namkeleja et al. (2013), this invasive species commonly outcompetes and displaces agricultural crops and native plant species. Its ability to invade is aided by allelopathy, a large seed bank, and resistance to extreme dryness and poor soil [25, 27, 31]. Also, a number of mechanisms i.e., contaminated soils, seed products, and crops, make it easy for its seeds to spread [23, 27]. Allelochemicals produced by *A. mexicana* have the potential to directly or indirectly suppress the germination and/ or growth of crops and native plants that compete with it in the area [24, 31]. The ability to suppress the germination or growth of neighbouring native plants or crops is referred to as allelopathy [18, 30]. *Argemone mexicana* is known to phenolic compounds i.e., benzoic acid, cinnamic acid ((d) (E)-3-Phenylprop-2-Enoic Acid), p-hydroxybenzoic acid (4-Hydroxybenzoic Acid), salicylic acid (2-Hydroxybenzoic Acid), and vanillic acid (4-Hydroxy-3-Methoxybenzoic Acid), which are responsible for allelopathic effects (Fig 2).

**Fig. 2.**
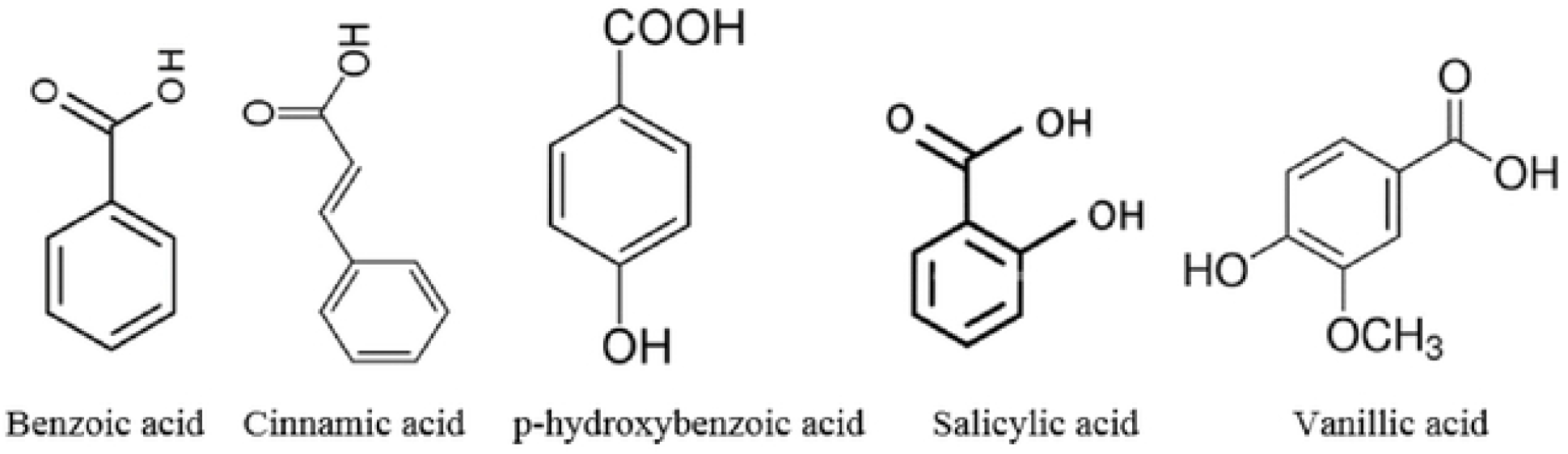
*Argemone mexicana’s* structures of phenolic allelochemicals [25, 27, 32]

*Argemone mexicana* is one of the deleterious invasive plants that has been increasingly invading natural and semi-natural habitats as well as agroecosystems in Tanzania. [29]. It has been invading grasslands and agricultural fields of maize (*Zea mays* L.), common bean (*Phaseolus vulgaris*), and other crops in the country [27, 29]. *Argemone mexicana* threaten the country’s ecosystem integrity, native biodiversity, economy, and food production [27]. Since it is toxic and the majority of grazers avoid it, it poses a risk to cattle and wildlife [27]. Its prickles could cause biodiversity and agricultural loss, as well as a decline in rangeland and/or grazing land quality [4]. They also annoy smallholder farmers and grazing animals [23, 25]. Health risks have also been linked to *A. mexicana* [25, 28] as well as suppressive effects on germination and growth of plants and crops [30, 32, 33]. For instance, A. mexicana was deemed a noxious IAP in South Africa due to the fact that when consumed, its seeds are detrimental for both human and animal health [23, 25, 28]. Cinnamic and benzoic acids are two examples of harmful allelochemicals that have been linked to a decrease in seed germination and seedling growth vigor in *A. mexicana* [25, 32, 34].

According to previous studies, the germination and seedling growth of various crops have been suppressed by the allelopathic effects of A. mexicana. Examples of these crops include tomato (*Solanum lycopersicum*), finger millet (*Eleusine coracana*), and cucumber (*Cucumis sativus*) [30, 32, 34, 35]. However, according to the available research, no study has been done in Tanzania to evaluate *A. mexicana’s* possible suppressive effects on legume crops, particularly *P. vulgaris* and *Z. mays*. Therefore, the study was carried out in petri dishes and pot experiments to investigate the suppressive effects of *A. mexicana* on the germination and seedling growth of *P. vulgaris* and *Z. mays. Argemone mexicana* leaf (AmL) crude extracts were used. It was hypothesized that AmL crude extract concentrations will negatively affect (i) seed germination and (ii) seedling stem height, stem diameter, root length, and fresh biomass of *P. vulgaris* and *Z. mays*. In general, this study is vital and intends to catalyze research on biological invasion to investigate the deleterious effects of IAPs across the world.

## Materials and methods

### Argemone mexicana leaf (AmL) crude extract

Fresh leaves of *A. mexicana* were collected from areas at Mbeya University of Science and Technology (MUST) (8° 56.24′ S and 33° 25.04′ E, 1636 m a.s.l.), farms (8° 56.45′ S and 33° 25.40′ E, 1643 m a.s.l.), and Iyunga (8° 55.85′ S, 33° 25.05′ E, 1616 m a.s.l.) in the Mbeya region between May and June 2022. The leaves were collected in the morning between 6:00 a.m. and 7:30 a.m. to avoid the probable degradation of non–photostable allelochemicals by the sun [18]. Collected AmL samples were kept in plastic paper bags and transported to the MUST biology laboratory (8°56.56′ S and 33°25.21′ E, 1651 m a.s.l.) for processing. They were cleaned with water to remove soil and/or debris particles. Afterwards, the AmL samples were air dried indoors at room temperature to avoid possible degradation of allelochemicals by ultraviolet (UV) light [18]. The dried AmL were ground into powder and stored in porous paper envelopes. About 100 g of AmL powder were measured using a digital balance and soaked in 1 l of distilled water. The crude was stored in a 4 l plastic container for 48 h in a dark room, and subsequently, the crude extract was filtered using muslin cloth. To get different aqueous concentrations, i.e., 0%, 25%, 50%, 75%, and 100% (w/v) of AmL (100 ml each), relative to the original extract, the filtrates were diluted with distilled water [18, 36]. The preparation procedures for crude extract concentrations followed those described in Ojija (2021) and Salih et al. (2021). The number of seed germinated, seedling stem height, stem diameter, root length, and fresh biomass, were used as indicators of *A. mexicana* suppressive effects on *P. vulgaris* and *Z. mays* (Nxumalo et al. (2022).

### Germination experiment

The *P. vulgaris* and *Z. mays* seeds were purchased from Ikuti market (S8° 56.07′, E33° 25.18′, 1630 m) in Mbeya region. To investigate the suppressive effect of AmL crude extracts on *P. vulgaris* seed germination, petri dish experiments were conducted at the MUST biology laboratory (S8° 56.56′, E33° 25.21′, 1651 m). Five glass petri dishes (each with a 70.84 cm^2^ surface area) per treatment were used and then replicated five times to make 50 petri dishes, i.e., 25 *P. vulgaris* and 25 for *Z. mays*. Petri dishes were rinsed with distilled water, dried, and lined with absorbent cotton wool before sowing 15 seeds of *P. vulgaris* and *Z. mays* in each petri dish. The seeds were irrigated ad libitum (i.e., kept moist) with different concentrations, i.e., 0%, 25%, 50%, 75%, and 100% (w/v) of AmL. The number of seeds that germinated was recorded daily for 16 days. The 16-day petri dish experiment was within the maximum germination period of *P. vulgaris* and *Z. mays* seed, which ranges between 7 and 12 days [37–39]. The positions of the petri dishes were randomized three times per week to ensure that sunlight was distributed similarly throughout both investigations. The criteria used for seed germination during the experiment was the emergence of the radicle [30]. The number of seed germinated was calculated and compared between AmL crude concentrations.

### Seedling growth experiment

Seedling growth experiments were conducted at MUST in a screen house (8° 56.61′ S and 33° 25.05′ E, 1646 m a.s.l.). The screen house protected the seedlings from damaging insects i.e., aphids and white flies. Fifteen seeds (15) of *P. vulgaris* and *Z. mays* were sowed in twenty–five pots (2 l) each. Pots were watered thoroughly at the time of sowing (i.e., 0.5 l per pot). Following five days of germination, *P. vulgaris* and *Z. mays* seedlings were irrigated three times per week with different AmL crude concentrations of 0%, 25%, 50%, 75%, and 100% (w/v). In addition to irrigation, the seedlings were also sprayed ad libitum with AmL concentrations twice per week using a hand sprayer. The seedlings were treated with AmL crude concentrations for 20 days between June and July 2022. The positions of the pots were randomized three times per week to ensure that sunlight was distributed similarly throughout both investigations. Following the experiment, total fresh biomass, root lengths, stem diameters, and stem heights were measured in order to examine the suppressive effects of AmL crude concentrations on *P. vulgaris* and *Z. mays* seedling growth. An analytical digital balance was used to quantify the seedlings’ total fresh biomass; digital callipers were used to measure the diameter of the stems above the first two leaves; and a meter ruler was used to measure the seedlings’ heights and root lengths.

### Statistical data analysis

The number of seed germinated and growth parameters (fresh biomass, root lengths, stem diameters, and heights) *P. vulgaris* and *Z. mays* were compared for different AmL crude extract concentrations using a one–way ANOVA and Kruskal–Wallis for parametric and non–parametric data, respectively. Levene’s test and Shapiro–Wilk test were used to test for equal variance and normality for all data, respectively. When the parametric assumptions were not confirmed after transformations (using Box-cox and/ or log transformation), the non–parametric Kruskal–Wallis test was used. Significant differences were confirmed using the post hoc Tukey–Kramer HSD and Mann–Whitney pairwise comparison tests. A 0.05 significance level was used for all the tests. Statistical tests were performed with Origin version 9.0 SR1 (2013).

## Results

### Seed germination under treatments

The results show that both *P. vulgaris* and *Z. mays* germination was suppressed by the AmL crude extracts at higher concentrations (75% and 100%, Fig. 4). The number of *P. vulgaris* and *Z. mays* seeds germinated under these concentrations was fewer compared to those germinated at lower concentrations (25% and 50%) and control (0%). The result indicates further that *P. vulgaris* and *Z. mays* seed germination decreased with increasing AmL crude extract concentrations (Fig. 4). For instance, the number of *Z. mays* seeds germinated at 25% and 50% is approximately four times that germinated at 100% (Fig.4). Overall, the number of seeds germinated at lower concentrations (i.e., 25% and 50%) and in the control differed from those germinated at higher concentrations (*P. vulgaris*: F_(4,20)_ = 5.28, p = 0.0046, Z. mays: F_(4,20)_ = 21.57, p < 0.0001, Fig 4).

**Fig. 3.**
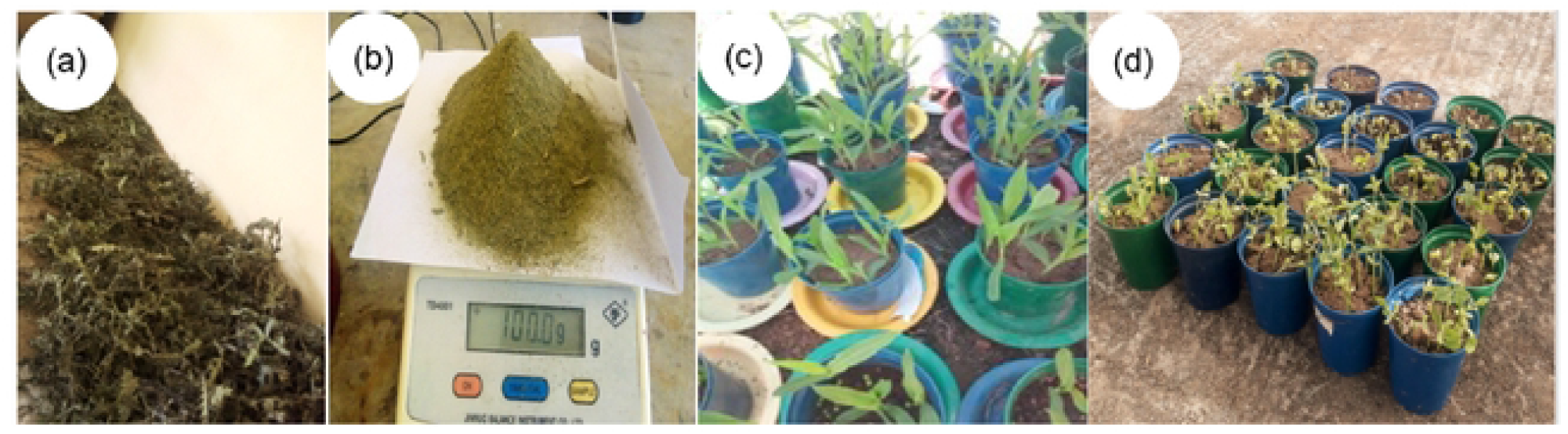
Pictures show the (a) dried leaves and (b) ground leaves of an invasive *Argemone mexicana*, and (c) *Zea mays* and (d) *P. vulgaris* seedlings. Photos: F. Ojija, 2022.

**Fig. 4.**
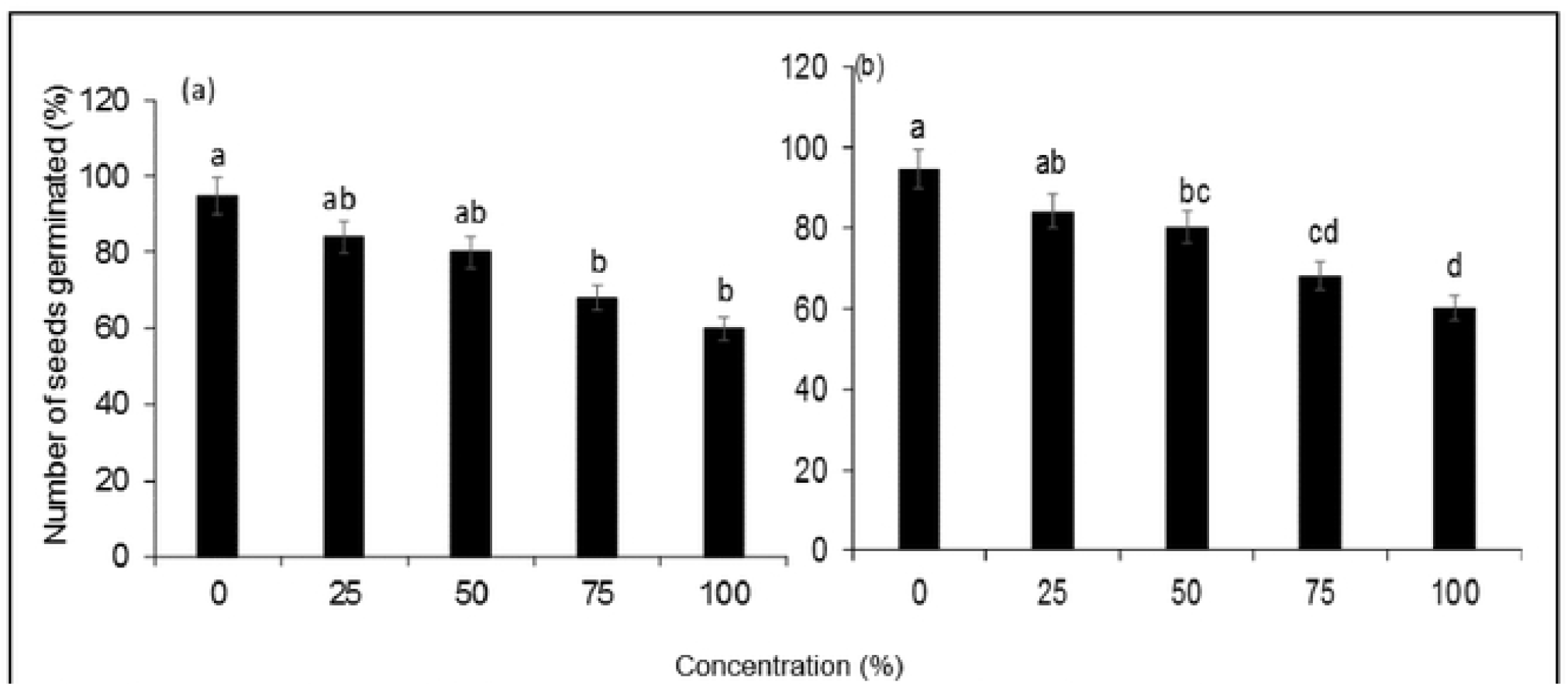
Mean number of (a) *P. vulgaris* and (b) *Z. mays* seed (± SE) germinated under different concentrations of AmL crude extracts over a 16-day experiment in petri dishes. The germination of *P. vulgaris* and *Z. mays* seeds decreased with increasing *A. mexicana* leaf crude concentrations. Bars with different letter (s) are significantly different at p = 0.05.

### Seedling growth parameters under treatments

Seedlings growth vigour of *P. vulgaris* and *Z. mays* was negatively affected since their growth parameters (stem diameters, stem heights, fresh biomass, and root lengths) under 75% and 100% AmL crude concentrations were lower (Fig 5, Table 1). The growth parameters of both *A. vulgaris* and *Z. mays* showed a significant difference between concentrations (Table 1). Fig 5 reveals a significant decrease in growth parameters of both test plants as treated with AmL crude extract. Stem height (Mean ± SE) of *P. vulgaris* and *Z. mays* seedlings treated with AmL crude extract differed significantly across different concentrations (Table 1). For instance, the stem height of *P. vulgaris* at 100% higher concentrations was reduced by 9.7 ± 0.1 cm, 7.5 ± 0.0 cm, and 5.2 ± 0.0 cm compared to 0%, 25%, and 50% concentrations, respectively (Fig 5, Table 1). Further, at 100% concentrations, the stem height of *Z. mays* was reduced by 6.9 ± 0.0 cm, 5.8 ± 0.8 cm, and 3.8 ± 0.0 cm compared to 25%, 50%, and 0%, respectively (Fig 5, Table 1). Similarly, the mean (± SE) stem height of both *P. vulgaris* and *Z. mays* seedlings treated with 75% concentrations was shorter than those grown at lower crude extract concentrations and controls (Fig 5, Table 1). For instance, at 75% crude extract concentration, *Z. mays* seedlings stem height was 2.0 ± 0.0 cm shorter or more compared to seedlings treated with lower concentrations and controls (Fig 5, Table 1).

**Fig. 5.**
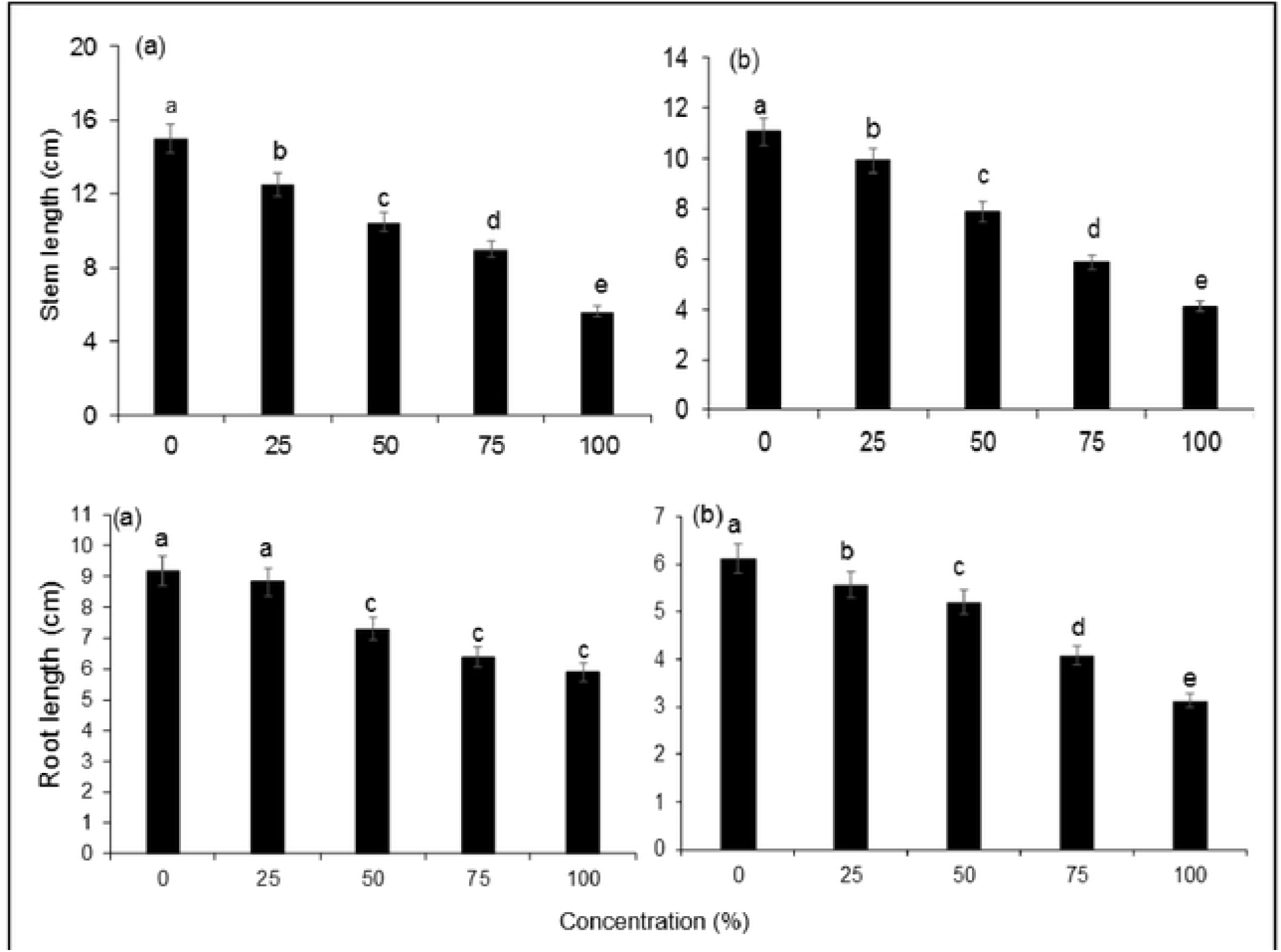
Mean growth parameters of (a) *P. vulgaris* and (b) *Z. mays* (± SE) seedlings treated with different concentrations of *A. mexicana* crude extract for 20 days in pot experiments. Bars with different letter (s) are significantly different at p = 0.05.

**Table 1.**
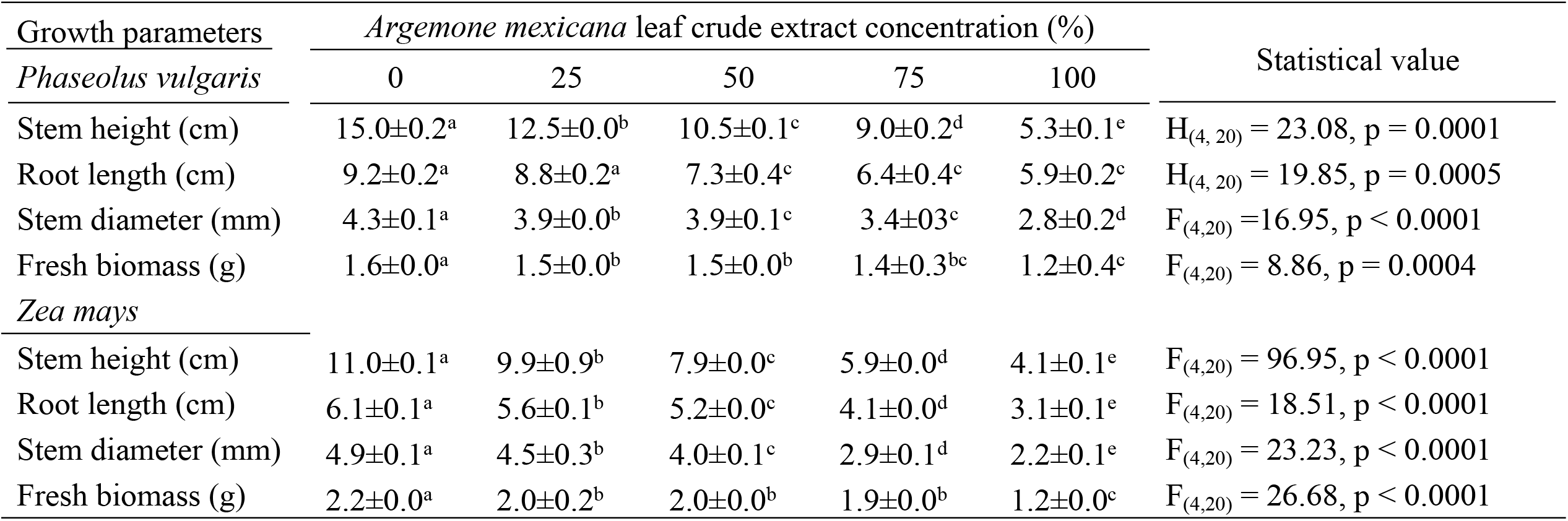
Kruskal–Wallis rank sum and one–way ANOVA test of *P. vulgaris* and *Z. mays* seedling parameters after 20 days of treatment in a screen house experiment. Values with different letter(s) in a row are significantly different by Tukey-Kramer HSD and Mann–Whitney pairwise comparison tests at p = 0.05.

| Growth parameters | *Argemone mexicana* leaf crude extract concentration (%) |  |  |  |  | Statistical value |
| --- | --- | --- | --- | --- | --- | --- |
| *Phaseolus vulgaris* | 0 | 25 | 50 | 75 | 100 |  |
| Stem height (cm) | 15.0±0.2<sup>a</sup> | 12.5±0.0<sup>b</sup> | 10.5±0.1<sup>c</sup> | 9.0±0.2<sup>d</sup> | 5.3±0.1<sup>e</sup> | $H_{(4, 20)} = 23.08, p = 0.0001$ |
| Root length (cm) | 9.2±0.2<sup>a</sup> | 8.8±0.2<sup>a</sup> | 7.3±0.4<sup>c</sup> | 6.4±0.4<sup>c</sup> | 5.9±0.2<sup>c</sup> | $H_{(4, 20)} = 19.85, p = 0.0005$ |
| Stem diameter (mm) | 4.3±0.1<sup>a</sup> | 3.9±0.0<sup>b</sup> | 3.9±0.1<sup>c</sup> | 3.4±0.3<sup>c</sup> | 2.8±0.2<sup>d</sup> | $F_{(4,20)} = 16.95, p < 0.0001$ |
| Fresh biomass (g) | 1.6±0.0<sup>a</sup> | 1.5±0.0<sup>b</sup> | 1.5±0.0<sup>b</sup> | 1.4±0.3<sup>bc</sup> | 1.2±0.4<sup>c</sup> | $F_{(4,20)} = 8.86, p = 0.0004$ |
| *Zea mays* |  |  |  |  |  |  |
| Stem height (cm) | 11.0±0.1<sup>a</sup> | 9.9±0.9<sup>b</sup> | 7.9±0.0<sup>c</sup> | 5.9±0.0<sup>d</sup> | 4.1±0.1<sup>e</sup> | $F_{(4,20)} = 96.95, p < 0.0001$ |
| Root length (cm) | 6.1±0.1<sup>a</sup> | 5.6±0.1<sup>b</sup> | 5.2±0.0<sup>c</sup> | 4.1±0.0<sup>d</sup> | 3.1±0.1<sup>e</sup> | $F_{(4,20)} = 18.51, p < 0.0001$ |
| Stem diameter (mm) | 4.9±0.1<sup>a</sup> | 4.5±0.3<sup>b</sup> | 4.0±0.1<sup>c</sup> | 2.9±0.1<sup>d</sup> | 2.2±0.1<sup>e</sup> | $F_{(4,20)} = 23.23, p < 0.0001$ |
| Fresh biomass (g) | 2.2±0.0<sup>a</sup> | 2.0±0.2<sup>b</sup> | 2.0±0.0<sup>b</sup> | 1.9±0.0<sup>b</sup> | 1.2±0.0<sup>c</sup> | $F_{(4,20)} = 26.68, p < 0.0001$ |

The stem diameter (Mean ± SE) of *P. vulgaris* and *Z. mays* seedlings differed significantly under different AmL crude extract concentrations (Fig. 6, Table 1). The diameter of seedlings treated with 100% crude concentrations was slightly smaller than that of those treated with 0%, 25%, and 50% crude extract concentrations (Fig. 6, Table 1). Moreover, the fresh biomass (Mean ± SE) of *P. vulgaris* and *Z. mays* seedlings was reduced at higher concentrations of AmL (Fig. 6, Table 1). For instance, the fresh biomass of seedlings from both test plants treated with 100% and 75 crude concentrations was slightly smaller than that of those treated with 0%, 25%, and 50% crude extract concentrations (Fig. 6, Table 1).

**Fig. 6.**
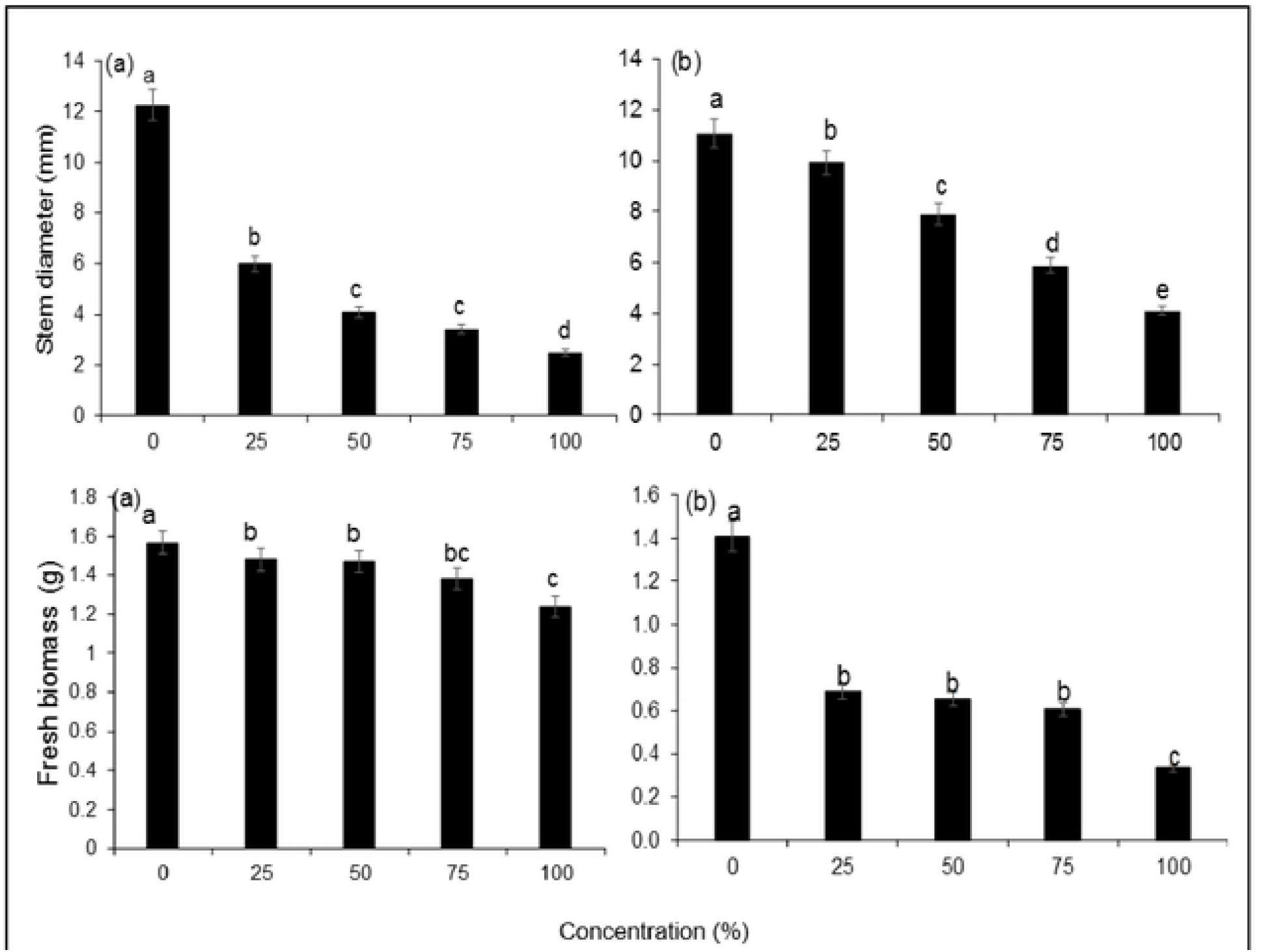
Mean growth parameters of (a) *P. vulgaris* and (b) *Z. mays* (± SE) treated with different concentrations of *A. mexicana* (AmL) crude extract for 20 days in pot experiments. Bars with different letter (s) are significantly different at p = 0.05.

## Discussion

The study investigated the phytotoxic effects of *A. mexicana* crude extract on the germination and growth of plants. Evidence was found for negative allelopathic effects of AmL crude extracts on *P. vulgaris* and *Z. mays* seed germination and growth. The results further demonstrated that germination and seedling growth of *P. vulgaris* and *Z. mays* were strongly suppressed at high AmL crude concentrations. This indicates that the effectiveness of AmL crude extracts is dosage–dependent, as supported by the previous studies [18, 36, 40, 41]. The AmL crude extract negatively affected *P. vulgaris* and *Z. mays* growth vigour as evidenced by reduced stem height, stem diameter, root length, and fresh biomass at high concentrations. This is further revealed by the number of *P. vulgaris* and *Z. mays* seeds that germinated under high AmL crude concentrations. The findings are consistent with earlier studies that found that *A. mexicana* can inhibit the germination and growth of plants and crops [24, 27, 30, 33, 34, 42].

For instance, Salih et al. (2021) found that *A. mexicana* extract negatively reduced seed germination, biomass, stem height, and root length of *Corchorus olitorus* and *Cassia senna* particularly at high concentrations. Similarly, [27] established that AmL and seed extracts suppressed seed germination, biomass, shoot height, and root length of *Brachiaria dictyoneura* L and Clitoria *ternatea* L seedlings. Conversely, other studies also claimed that *A. mexicana* possesses allelochemicals that are responsible for suppressive or negative allelopathic effects on other plants. Thus, the current study indicates the allelochemicals present in AmL might be responsible for inhibiting seed germination and growth of *P. vulgaris* and *Z. mays*. These allelochemicals have the ability to interfere with the physiological mechanisms involved in the germination and growth of plant species [18, 43–45].

Some of the allelochemicals reported from previous studies to be responsible for negative allelopathic effects include cinnamic acid, vanillic acid, benzoic acid, salicylic acid, and p-hydroxybenzoic acid [24, 25, 29]. It has been stated that *A. mexicana* uses p-hydroxybenzoic and vanillic acids to interfere with the water balance of native plants and crops and subsequently suppress their growth [25, 32]. Because of this, the physiological characteristics of adjacent plants are negatively affected, which in turn inhibits root activity, reduces the amount of chlorophyll, and then increases the rate of photosynthesis [18, 32]. Also, [34] reported that the growth and germination of soybean were inhibited following treatment with high concentrations of p-hydroxybenzoic and vanillic acids. Also, [46] made similar observation that vanilic acid was found to suppress eggplant (*Solanum melongena*) seed germination and seedling growth. This shows that allelochemicals (e.g., p-hydroxybenzoic, salicylic, and vanillic acids) present in AmL might be responsible for the suppressive effects observed on the germination and growth of *P. vulgaris* and *Z. mays*.

Additionally, the ability of *A. mexicana* to suppress *P. vulgaris* and *Z. mays* seed germination and growth in this study could be due to salicylic and cinnamic acids [24]. These allelochemicals were previously found to inhibit seed germination and seedling growth of cowpea (*Vigna unguiculata*) [42, 47], *S. melongena* [46], and *Z. mays* [24] at high concentrations. Therefore, these allelochemicals that are present in AmL can also negatively affect *P. vulgaris* and *Z. mays* germination, seedling growth, and fresh biomass, as found in this study [29]. Nevertheless, it should be understood that the effectiveness of allelochemicals present in an invasive plant can be species–specific. This means that some plant species may experience positive allelopathic effects while others may experience negative effects. For example, according to Burhan and Shaukat (1999) findings, the effects of phytotoxin from *A. mexicana* appeared to be species–specific, as not all of the studied species were equally suppressed by the extract. Generally, *A. mexicana* has the potential to suppress the germination and early–growth of plants, including crops, as evidenced from the current and previous studies.

## Conclusions

Since the invasive *A. mexicana* exhibits negative allelopathic effects on other plants in addition to being detrimental to humans and livestock, it should be controlled. But the involvement of local communities, especially farmers and pastoralists, in the management of the invasive species is vital because they are directly affected by *A. mexicana*. Besides, this study’s results showed that *A. mexicana* crude extract reduced *P. vulgaris* germination and growth; nonetheless, field research is required to fully understand the allelopathic effects of *A. mexicana* on other species of plants.

## Acknowledgements

The author is greatly acknowledging the valuable in–kind support from Mbeya University of Sciences (MUST) and the Social Health and Environment Management Organization (SHEMO). He is also thankful to Leticia P. Lutambi, Tareq S. Said, Thobias Protaz, Rachel Palangyo, Lusekelo S. Adam, Eclan G. Raphael, and Dorine C. Mkwawa for assisting with the field and laboratory work.

